# Resolving replication incompatibility between chloroplast and conjugative plasmids in *E. coli*

**DOI:** 10.1101/2025.10.16.682956

**Authors:** Emma Jane Lougheed Walker, Tahani Jaafar, Ayagiysan Kaneshan, Bogumil Jacek Karas

**Author notes:** The author responsible for distribution of materials integral to the findings presented in this article in accordance with the policy described in the Instructions for Authors is Bogumil Karas.

## Abstract

Chloroplast genomes present a promising chassis for engineering photosynthetic eukaryotes, but efficient delivery of large DNA constructs back into the organelle remains a major technical barrier. Conventional transformation methods rely on purified DNA and physical force to drive uptake into the chloroplast, often resulting in DNA shearing and thus low transfer efficiency for large constructs. Bacterial conjugation offers an attractive alternative as this is an entirely *in vivo* process, enabling DNA transfer without any physical manipulation. To assess the feasibility of this approach, we attempted to generate an *Escherichia coli* donor strain carrying both the broad-host-range conjugative plasmid pTA-Mob (52.7 kb) and the cloned *Phaeodactylum tricornutum* chloroplast genome (pPt_Cp, 132.9 kb). Unexpectedly, pTA-Mob and pPt_Cp proved incompatible: co-maintenance could not be achieved via electroporation, and conjugation with a self-transmissible pTA-Mob variant resulted in a ∼10□-fold decrease in transfer efficiency. Systematic testing of pTA-Mob and pPt_Cp plasmid variants as well as sequence analysis of evolved transconjugants revealed that the incompatibility arose from the pBBR1 replicon present in pTA-Mob. Guided by these insights, we identified an alternative conjugative plasmid, pRL443, that was compatible with pPt_Cp. Together, these findings provide a framework for dissecting plasmid incompatibility when working with large constructs and establish a functional conjugative system capable of mobilizing the *P. tricornutum* chloroplast genome.

## INTRODUCTION

Photosynthesis is a fundamental biological process that sustains virtually all life on Earth. This process first originated in an ancient cyanobacterial lineage over 3 billion years ago and eventually led to the oxygenation of our oceans, soils, and atmosphere – a geobiological event that paved the way for the evolution of complex life forms and ecosystems (Lyons et al., 2014; Sánchez-Baracaldo and Cardona, 2020). Eukaryotes are thought to have acquired photosynthesis nearly 2 billion years later by forming an endosymbiotic relationship with an ancient cyanobacterial species (Sibbald and Archibald, 2020). This is in part evidenced by the fact that all extant photosynthetic eukaryotes possess a light-harvesting organelle (i.e., chloroplast) that harbours its own highly reduced prokaryotic-like genome, akin to the mitochondria and its organellar genome.

Photosynthetic organisms, whether of prokaryotic or eukaryotic origin, once again hold immense potential for the future of this planet by virtue of their ability to sequester and convert atmospheric carbon dioxide into useful organic compounds. If leveraged accordingly, photosynthesis could be used as a tool to mitigate climate change and to drive sustainable biofactories. Unlocking the full potential of photosynthesis, however, will require the development of robust large-scale engineering tools for these organisms.

Chloroplast genomes offer a unique chassis for high-throughput engineering of photosynthesis. These genomes typically range in size from 100 to 200 kb and encode for 100 to 250 genes (Lang and Nedelcu, 2012), making them orders of magnitude smaller than the average eukaryotic or prokaryotic chromosome. These reduced genomes are also considered to be “naked”, not bearing any histones or other DNA-binding proteins that could otherwise influence transcription (Bock, 2015). Taken together, these facets make the design, synthesis and assembly of designer chloroplast genomes attainable; indeed, the chloroplast genomes of rice (*Oryzae sativa*, 135 kb; Itaya et al., 2008), *Chlamydomonas reinhardtii* (205 kb; O’Neill et al., 2012), and *Phaeodactylum tricornutum* (117 kb; Walker et al., 2024) have been reconstructed exogenously from large DNA fragments before. These assembly pipelines offer the utmost control over genome design, making it possible to conceive of and create chloroplast genomes that are minimized, recoded, refactored, and more (Walker et al., 2024). *De novo* genome assembly is only one part of the design-build-test cycle for creating synthetic chloroplast though; to develop a chloroplast controlled by a synthetic genome, effective methods for delivering and installing partial or complete chloroplast genomes are also needed.

Few methods have been explored for large-scale transformation of the chloroplast genome. The cloned *C. reinhardtii* chloroplast genome was delivered to the chloroplast through biolistic particle bombardment (O’Neill et al., 2012), yet ultimately could not be successfully installed (i.e., achieve homoplasmy). This was hypothesized to be due to recombination between the engineered and endogenous genomes but could have also been influenced by DNA shearing prior to installation, as the 242 kb cloned genome would be subject to shear forces during isolation, preparation for delivery, and biolistic bombardment itself. In the time since, biolistics has been used to deliver and successfully install recoded portions of the *C. reinhardtii* chloroplast genome, with regions as large as 9.1 kb being replaced by synthetic counterparts (Mordaka et al., 2025). This demonstrates that biolistics could be used for stepwise replacement of the chloroplast genome in lieu of whole genome replacement and presents a strategy to achieve homoplasmy, as extensive recoding prevents homologous recombination between the endogenous and engineered genomes (Mordaka et al., 2025). Delivery of the *P. tricornutum* chloroplast genome was not attempted as reliable selective markers and an approach to prevent unwanted homologous recombination had yet to be established (Walker et al., 2024); however, the use of bacterial conjugation was proposed as an alternative delivery method to biolistics. To facilitate this, an origin of transfer (*oriT*) sequence was included in the cloning vector used during assembly of the *P. tricornutum* chloroplast genome (Walker et al., 2024).

Bacterial conjugation holds immense potential for intact delivery of whole chloroplast genomes. Here, a bacterial donor strain harbours a conjugative plasmid that encodes for a pilus and type IV secretion system, along with a mobilizable plasmid bearing an *oriT* sequence (Costa et al., 2024). The *oriT* is recognized and covalently bound by relaxase, a conjugative protein that is involved in the physical transfer of the plasmid between the donor and recipient cell (Costa et al., 2024). In some cases, the conjugative plasmid itself can also carry an o*riT*, making it self-transmissible. Once conjugation is initiated, the mobilizable plasmid is transferred from the donor bacterium to a recipient cell, which may be prokaryotic or eukaryotic depending on the host range of the conjugative plasmid (Karas et al., 2015; Soltysiak et al., 2019). It is unknown whether plasmid transfer takes place through the lumen of the pilus or via cell-to-cell contact (Costa et al., 2024), but in any case, DNA transfer occurs *in vivo*, thereby avoiding shear forces and other forms of extracellular DNA damage. Relaxase is thought to mediate the import of the protein–DNA complex into the host cell nucleus during conjugation to a eukaryotic recipient (Silby et al., 2007; Tinland et al., 1992), leading to high-efficiency transfer rates upon successful cell-to-cell contact. Conjugation has been shown to intactly transfer plasmids upwards of 1.6 Mb before (Blanca-Ordóñez et al., 2010), demonstrating its propensity for moving constructs far larger than the average chloroplast genome.

While the chloroplast genomes of *C. reinhardtii* and *P. tricornutum* were able to be cloned and stably maintained in *E. coli* (O’Neill et al., 2012; Walker et al., 2024), their potential for transfer via bacterial conjugation was not investigated in the previous studies. From a size perspective, such transfer should be feasible; however, chloroplast genomes contain prokaryotic replication origins and other elements required for stability and partitioning. If these regions share sequence similarity to those present in a conjugative plasmid, competition for replication and maintenance machinery can occur, leading to plasmid incompatibility (Novick, 1987). This ultimately results in the partial or complete loss of one of the plasmids over time (Novick, 1987). Therefore, the compatibility of the chloroplast genome with broad-host-range conjugative plasmids must be assessed before conjugation can be proposed as a viable delivery mechanism for chloroplast engineering.

In this paper, we investigate bacterial conjugation as a means to transfer the cloned *Phaeodactylum tricornutum* chloroplast genome between cells. We found that the cloned chloroplast genome, pPt_Cp, is incompatible with the broad-host-range plasmid pTAMob. Through sequence analysis of evolved transconjugants and systematic testing of pTAMob deletion variants, we identified the genetic determinants underlying this incompatibility on both the conjugative plasmid and the chloroplast genome. Building on this insight, we characterized an alternative broad-host-range plasmid, pRL443, which proved compatible with the cloned chloroplast genome. Together, these findings establish first demonstration of conjugation being used to mobilize the whole chloroplast genome, providing a foundational framework for future genome-scale engineering of photosynthetic organelles.

## RESULTS

### Conjugating pTAMob into pPt_Cp and partial-genome strains

To begin our studies, we attempted to electroporate pTAMob (52.7 kb), a broad host range RK2-based conjugative plasmid (Strand et al., 2014), into an *E. coli* EPI300 strain harbouring pPt_Cp (132.9 kb, Addgene ID: 206855), the cloned *P. tricornutum* chloroplast genome. Multiple attempts did not yield any transformants, despite being able to electroporate both pTAMob and pPt_Cp individually into EPI300 cells with high efficiency (>1000 colonies per transformation). This suggested that an incompatibility between pTAMob and pPt_Cp was preventing successful maintenance of both plasmids in EPI300 simultaneously.

Conjugation is generally more efficient than electroporation for plasmid transfer in *E. coli*, particularly when transferring large or complex constructs. To further explore the hypothesized plasmid incompatibility, we conjugated pTAMob 2.0 (56.5 kb, Addgene ID: 149662), an engineered version of pTAMob that is self-transmissible (Soltysiak et al., 2019), into *E. coli* EPI300 strains harbouring partial chloroplast genome constructs (Fig. 1A). These constructs were assembled using various combinations of the initial eight fragments used in whole genome assembly (Walker et al., 2024).

**Figure 1.**
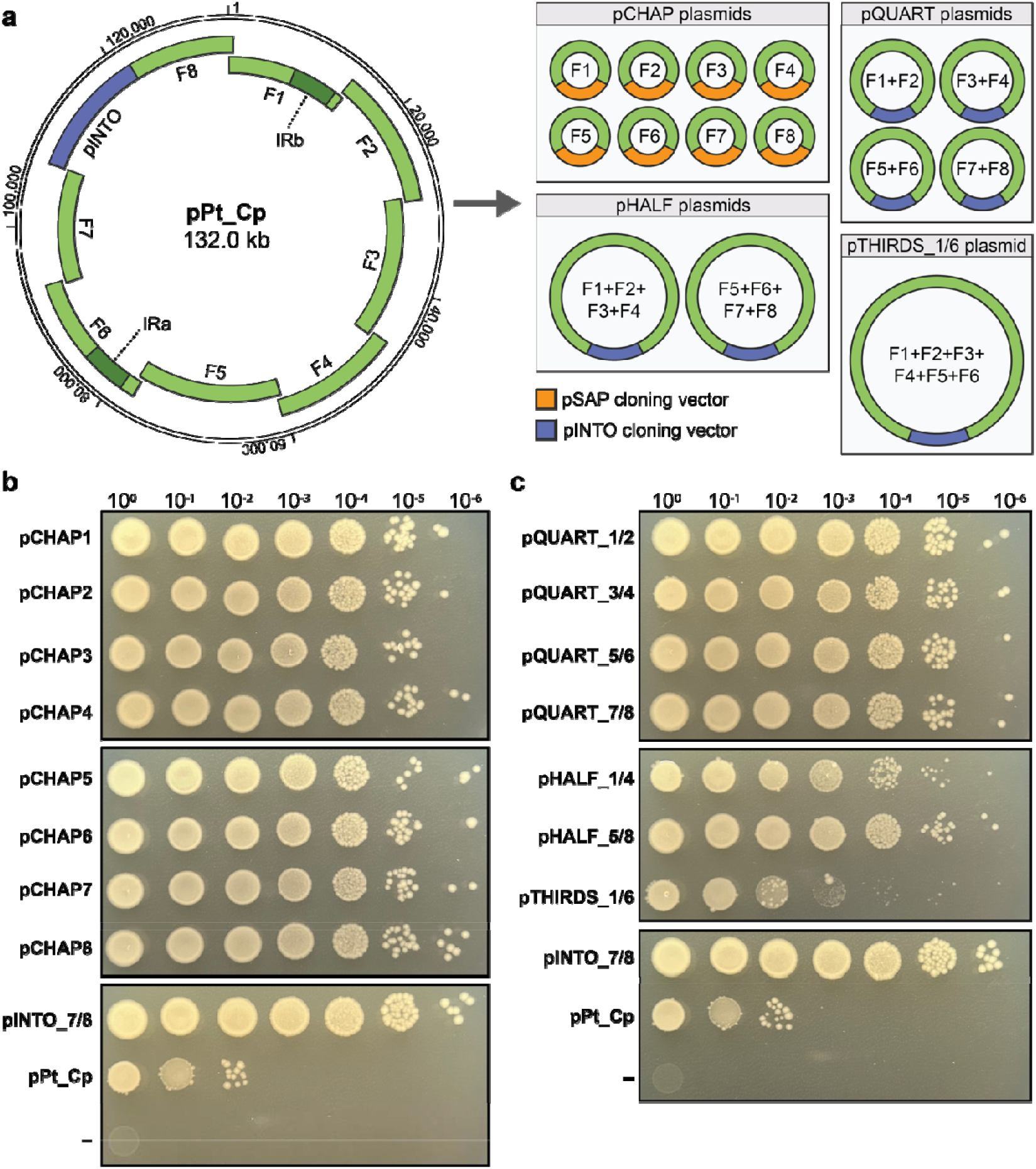
Conjugation of pTAMob 2.0 into *E. coli* strains containing partial chloroplast genomes. (**a**) The chloroplast genome was previously split into eight overlapping fragments and cloned as multiple partial genome variants. The pCHAP plasmids each harbour one-eighth of the genome, whereas the pQUART and pHALF plasmids each harbour a quarter or halve of the genome, respectively. The pTHIRDS plasmid harbours chloroplast genome fragments 1 to 6. The inverse repeat (**IR**) regions of the chloroplast genome are annotated on the pPt_Cp map. (**b**) Conjugation of pTAMob 2.0 into strains harbouring pCHAP1 to pCHAP8. These strains each carry a portion of the *P. tricornutum* chloroplast genome that ranges in size from 11.9 to 17.6 kb. (**c**) Conjugation of pTAMob 2.0 into strains harbouring a quarter (pQUART plasmids), half (pHALF plasmids), or three-quarters (pTHIRDS_1/6 plasmid) of the chloroplast genome. Strains carrying the whole chloroplast genome (i.e., pPt_Cp) or the cloning vector pINTO_7/8 were used as benchmarks for assessing conjugation efficiency. The negative control consists of the conjugative donor, pTAMob 2.0, by itself. Spot plating was performed on LB agar supplemented with chloramphenicol (15 µg/ml) and gentamycin (40 µg/ml).

The pCHAP plasmids each harbour an 11.9 to 17.6 kb region of the *P. tricornutum* chloroplast genome (Walker et al., 2024; Fig. 1A, Table S1). These regions span the entirety of the chloroplast genome and were previously captured with the pCC1BAC-based cloning vector pSAP (Walker et al., 2024). This cloning vector differs only slightly from pINTO_7/8, the backbone of pPt_Cp, in that it contains a different yeast selective marker (i.e., TRP1 instead of HIS3), cannot be mobilized (i.e., no *oriT*), and does not harbour a *P. tricornutum* selective marker.

The conjugation efficiency of pTAMob 2.0 into strains harbouring pCHAP1 through 8 did not differ from that of pINTO_7/8 (i.e., the positive control, Fig. 1B). When spot plated, single colony growth (SCG) was observable at dilutions of 10^-5^ and 10^-6^ for these conjugative crosses, whereas pPt_Cp only demonstrated SCG at a dilution of 10^-2^. This discrepancy in conjugation efficiency was even more apparent when the transconjugant cells were spread across entire plates (Fig. S1A), where a ∼10^5^ difference in SCG is visible between the pCHAP1 and pPt_Cp crosses. This conjugation experiment confirmed that pTA-Mob 2.0 and pPt_Cp are highly incompatible and revealed that this incompatibility does not arise from a single genomic region, but rather from interactions among multiple regions of the chloroplast genome.

To further explore the cause of incompatibility between pTAMob 2.0 and pPt_Cp, we assembled and performed conjugation to additional partial genome constructs. Using pINTO_7/8 as a cloning vector, the fragments originally captured in pCHAP1 through 8 were combined to form quartered, halved, and a three-quarters version of the chloroplast genome (Fig. 1A). These constructs were labelled as pQUART, pHALF, and pTHIRDS followed by a series of numbers that designates which chloroplast fragments were combined (e.g., pHALF_1/4 contains the genome fragments 1 to 4). Conjugation of pTAMob 2.0 into these strains demonstrated that incompatibility arises to some degree in the strains harbouring pHALF_1/4 and pTHIRDS_1/6 (Fig. 1C), though this drop in efficiency is not as extreme as with pPt_Cp. Like before, these discrepancies become more apparent when the transconjugant cells are spread across entire plates (Fig. S1B). This again indicates that incompatibility does not stem from a single genomic region, but rather from the combined effects of multiple regions, with the complete genome exhibiting the strongest incompatibility.

### Analysis of pTAMob 2.0 x pPt_Cp transconjugants

Conjugation of pTAMob 2.0 into the strain harbouring pPt_Cp generated big and small colony phenotypes (Fig. 2A). This pattern was also observed during conjugation of pTAMob 2.0 into the strains harbouring pHALF_1/4 and pTHIRDS_1/6 (Fig. S1B). Interestingly, big colonies could be successfully passaged for further analysis, whereas small colonies would not regrow upon passaging. This suggested that mutations to pTAMob 2.0 and/or pPt_Cp had spontaneously arisen in the big colonies, enabling the stable maintenance and propagation of both plasmids.

**Figure 2.**
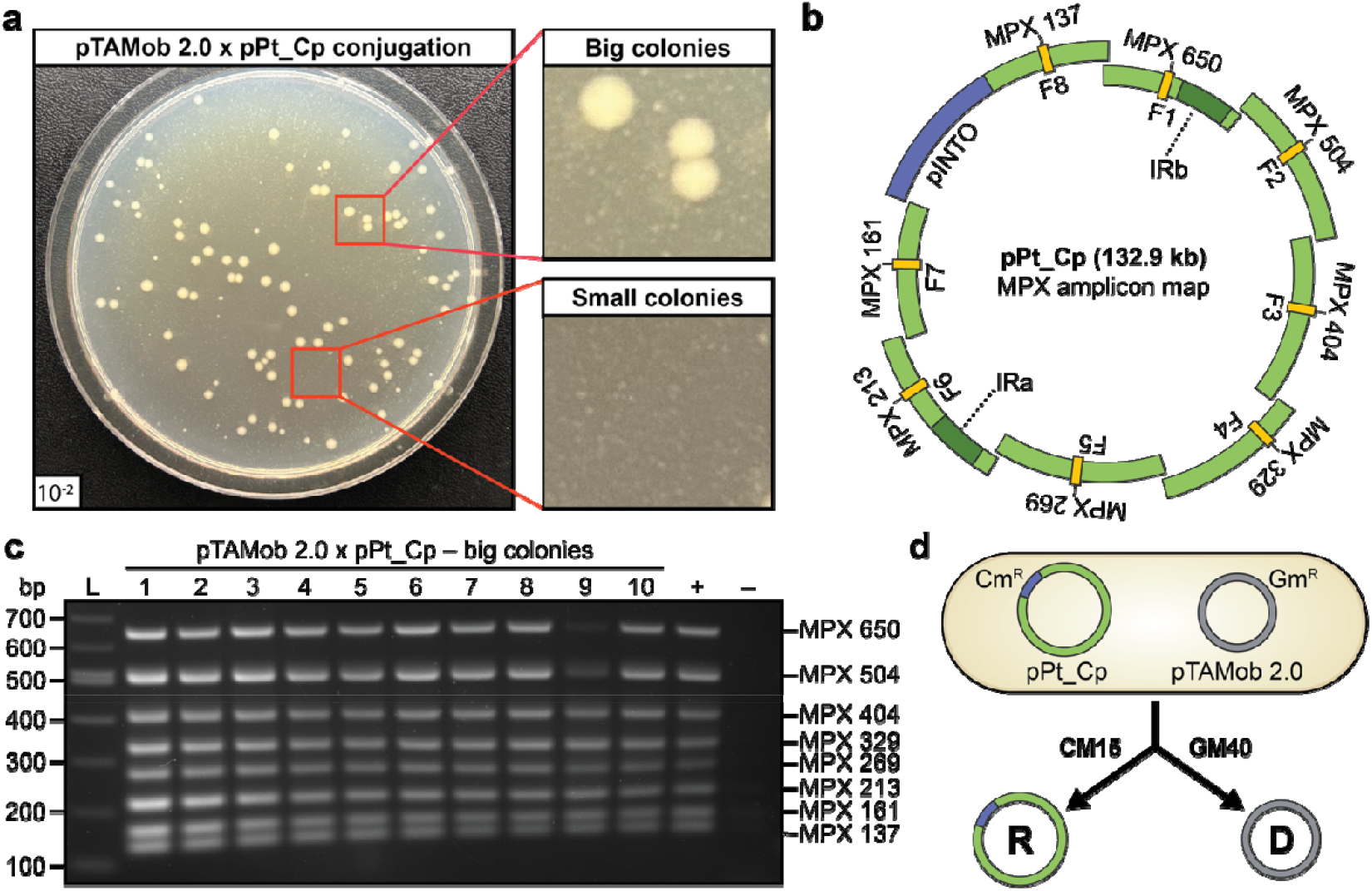
Analysis of pTAMob 2.0 x pPt_Cp transconjugant colonies. (**a**) Distinct big and small colonies are visible following conjugation of pTAMob 2.0 into pPt_Cp. (**b**) The cloned chloroplast genome, pPt_Cp, was originally captured as eight overlapping fragments (F1 to F8) using the cloning vector pINTO_7/8. MPX primers were positioned in the centre of these fragments to be able to screen pPt_Cp and the partial-genome variants used in this study. The inverted repeat (**IR**) regions A and B are also labelled on the plasmid map. (**c**) An MPX PCR screen of 10 big colonies following conjugation of pTAMob 2.0 into the strain harbouring pPt_Cp. (**d**) Isolation of pPt_Cp and pTAMob 2.0 from a transconjugant strain. Once isolated, the plasmids can be re-transformed into EPI300. Uptake of pPt_Cp will give resistance to chloramphenicol (CM15), whereas uptake of pTAMob 2.0 will give resistance to gentamycin (GM40). This was used to create new recipient (R) and donor (D) strains to test for putative mutations in pPt_Cp and pTAMob 2.0, respectively. These strains are denoted as Pt-C and Mob-C.

A multiplex (MPX) PCR screen spanning all eight fragments of the cloned chloroplast genome (Fig. 2B) was performed using DNA isolated from 10 big colonies to screen for any large rearrangements or deletions. All colonies demonstrated the expected MPX banding pattern, though colony 9 had noticeably weaker amplicons for fragments 1 and 2 (Fig. 2C). Plasmid DNA from the big colonies were then electroporated into EPI300 to recover individual strains carrying transconjugant pPt_Cp or pTAMob 2.0. (Fig. 2D). From each big colony, both a donor (i.e., pTAMob 2.0) and recipient (i.e., pPt_Cp) strain were generated to re-test conjugation and determine whether mutations in either plasmid had resolved the incompatibility.

Conjugation of the Mob-C1 to -C10 strains into the strain harbouring pPt_Cp showed no change in efficiency, indicating that pTAMob 2.0 had likely not undergone mutation in the big colonies (Fig. 3A). Conversely, conjugation of pTAMob 2.0 into the Pt-C1 to -C10 strains was ∼10^4^-fold more efficient than into the original pPt_Cp strain (Fig. 3B), indicating that mutations in the chloroplast genome of the big colonies had restored plasmid compatibility for all the tested strains. Relatedly, the colonies generated from the pTAMob 2.0 x Pt-C crosses no longer demonstrated the big and small phenotypes seen before (Fig. 2A); colonies were uniform in size and appearance, closely resembling those of the pINTO_7/8 control (Fig. 3B).

**Figure 3.**
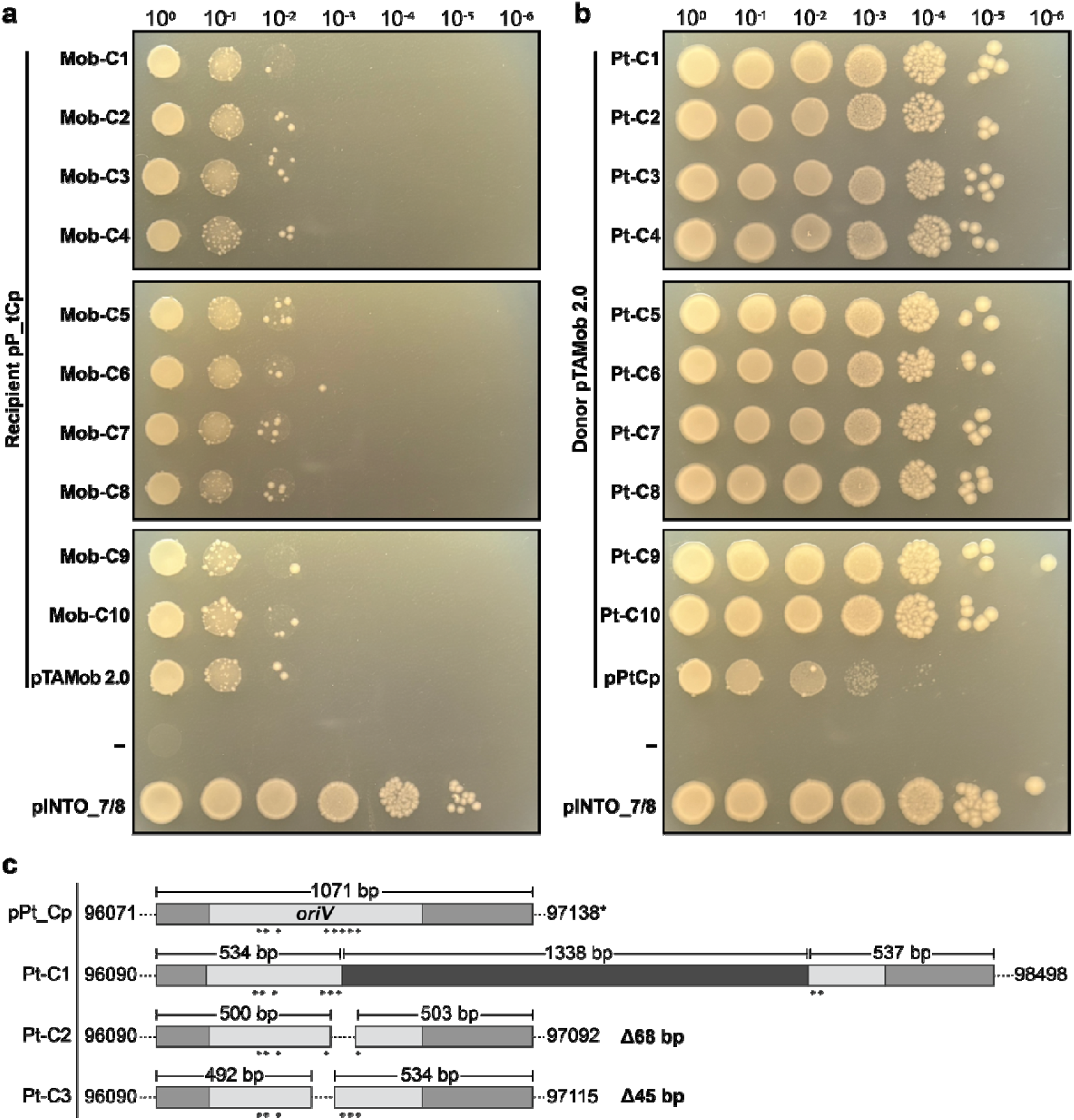
Conjugation using pTAMob 2.0 and pPt_Cp recovered from big colonies. (**a**) Conjugation of Mob-C1 to C-10 and the original pTAMob 2.0 into the strain harbouring pPt_Cp. The negative control consists of pPt_Cp by itself. (**b**) Conjugation of pTAMob 2.0 into Pt-C1 to C-10 and the original strain harbouring pPt_Cp. The negative control consists of pTAMob 2.0 by itself. The strain carrying the cloning vector pINTO_7/8 was used as a benchmark for assessing conjugation efficiency. Spot plating was performed on LB agar supplemented with chloramphenicol (15 µg/ml) and gentamycin (40 µg/ml). (**c**) Alignment of Pt-C1, C2, and C3 to pPt_Cp. All three colonies demonstrated indels in the *oriV* region, with Pt-C1 containing a 1338 bp insertion that maps to the *E. coli* genome. Iterons are indicated by arrows underneath the *oriV* sequence. The asterisk (*) denotes that 97,138 bp is the annotated endpoint of the *oriV* region on the sequenced pPt_Cp plasmid, reflecting a sequencing artefact; the actual coordinate should be 97,142 bp.

To identify mutations responsible for restored plasmid compatibility, DNA isolated from Pt-C1, -C2, and -C3 were sequenced and aligned to pPt_Cp (Fig. 3C). The sequences spanning the chloroplast genome (∼117 kb) remained unchanged across the analysed colonies; however, insertions and deletions (indels) were consistent in one region of the cloning vector. This region maps to the RK2 vegetative origin of replication (*oriV*) and results in disruption of its iteron sequences, which are tandem repeats necessary for initiation and regulation of plasmid replication (Maurya et al., 2023). Deletions in Pt-C2 and Pt-C3 remove three and two of the eight iterons, respectively, while the insertion in Pt-C1 interrupts the iteron repeats. When queried, the ∼1.3 kb insert exhibited 100% sequence identity to a region of the *E. coli* K-12 chromosome (GenBank accession: CP047127.1; positions 851261–852593), suggesting a sporadic chromosomal insertion event had taken place in this plasmid. Taken together, these findings demonstrate that disruption of the *oriV* sequence in pPt_Cp restores plasmid compatibility with pTAMob 2.0.

### Conjugation with minimized variants of pTAMob 2.0

To determine the source of incompatibility on pTAMob 2.0, we performed conjugation using minimized pTAMob derivatives generated from previous work (Fig. 4A; Cochrane et al., 2022). The deletions are non-essential to conjugation and span three regions comprising 24.7 kb and 41 open reading frames (ORFs, Table S2). All four minimized variants – M5C3, M6C4, M7C2, and M8C10 – were conjugated into the strain harbouring pPt_Cp (Fig. 4B, Fig. S2). Plasmids M7C2 and M8C10 restored conjugation efficiency, suggesting that the incompatibility lies within the MVC region, as deletion of the MV6 and MVA regions was not enough to restore efficiency.

**Figure 4.**
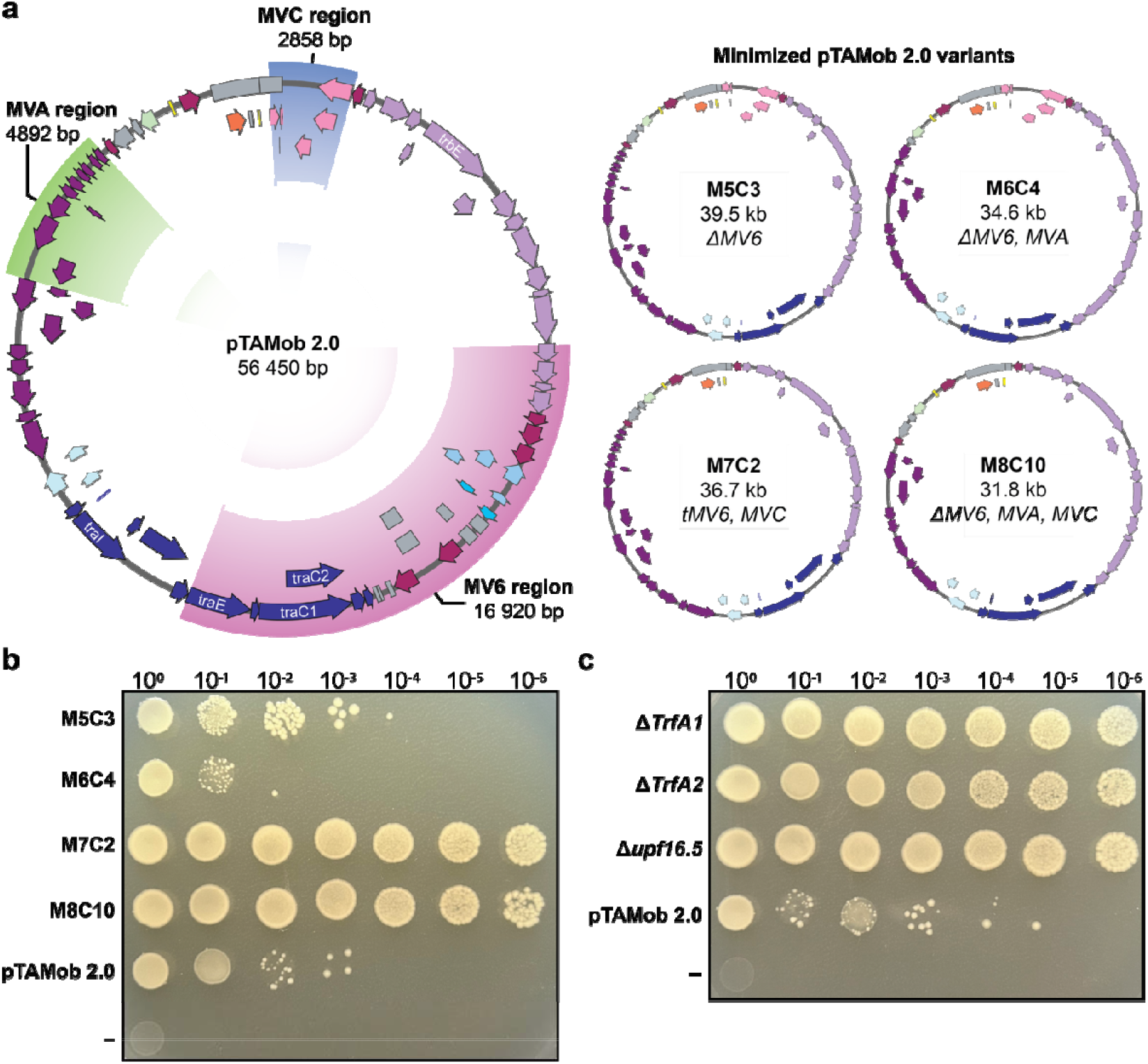
Conjugation using minimized or deletion derivatives of pTAMob 2.0. (**a**) A plasmid map of pTAMob 2.0 with the non-essential regions highlighted. MV6, MVA, and MVC were sequentially removed from pTAMob 2.0 to give rise to the minimized plasmids M5C3, M6C4, M7C2, and M8C10. The regions deleted are denoted underneath the plasmid names. (**b**) Conjugation of minimized and (**c**) single-gene deletion derivatives of pTAMob 2.0 into the strain harbouring pPt_Cp. The negative control consists of the original conjugative donor, pTAMob 2.0, by itself. Spot plating was performed on LB agar supplemented with chloramphenicol (15 µg/ml) and gentamycin (40 µg/ml).

The MVC region contains six ORFs, three of which are duplicated genes that were carried over when assembling the *oriT* into pTAMob 2.0 (Table S2; Soltysiak et al., 2019). The duplicated genes (i.e., *traJ, traX*, and *traI*) are derived from the Tra1 operon and are unlikely to be expressed as the regulatory elements (i.e., promoter, operator, terminator) are missing. The remaining ORFs encode for the replication initiator proteins TrfA1 and TrfA2, as well as upf16.5, which is functionally uncharacterized but may influence plasmid copy number (Bartosik et al., 2016; Stalker et al., 1981).

To further narrow the source of plasmid incompatibility, conjugation was performed using single-gene knock-out variants of pTAMob 2.0 that were previously generated (Fig. 4C; Cochrane et al., 2022). Knock-out of *trfA1, trfA2*, or *upf16*.*5* restored conjugation efficiency and thus resolved the plasmid incompatibility with pPt_Cp. These ORFs are contained within a putative operon, with *trfA2* also being nested inside the terminal region of *trfA1*, so the exact source of incompatibility cannot be elucidated any further without additional engineering of pTAMob 2.0. With that being said, it is known that TrfA is a replication initiator protein that recognizes and helps to establish replication at the *oriV* sequence (Kongsuwan et al., 2006). Given that *oriV* mutations restore compatibility in pPt_Cp, it is most likely that the TrfA1 and TrfA2 replication initiator proteins are responsible for the incompatibility observed with pTAMob 2.0.

### Conjugation using an alternative RK2-based plasmid

Our investigation into the source of plasmid incompatibility between pPt_Cp and pTAMob 2.0 suggested that replication of the *oriV* was leading to instability and plasmid loss. However, in past work, we have demonstrated that pPt_Cp can be induced to high-copy number replication at the *oriV* when maintained in the *E. coli* strain EPI300 (Walker et al., 2024), which has a genomic copy of *trfA* driven by the arabinose-inducible P_BAD_ promoter (Fig. 5A, first panel). High-copy number expression of pPt_Cp leads to a growth burden in *E. coli*, but does not cause any plasmid rearrangements or deletions that would be expected if *oriV*-based replication was unstable (Walker et al., 2024). Furthermore, all partial chloroplast genome constructs contained an *oriV* sequence within their cloning vector backbones and plasmid incompatibility was observed only in a subset of constructs – specifically those containing at least half of the chloroplast genome.

**Figure 5.**
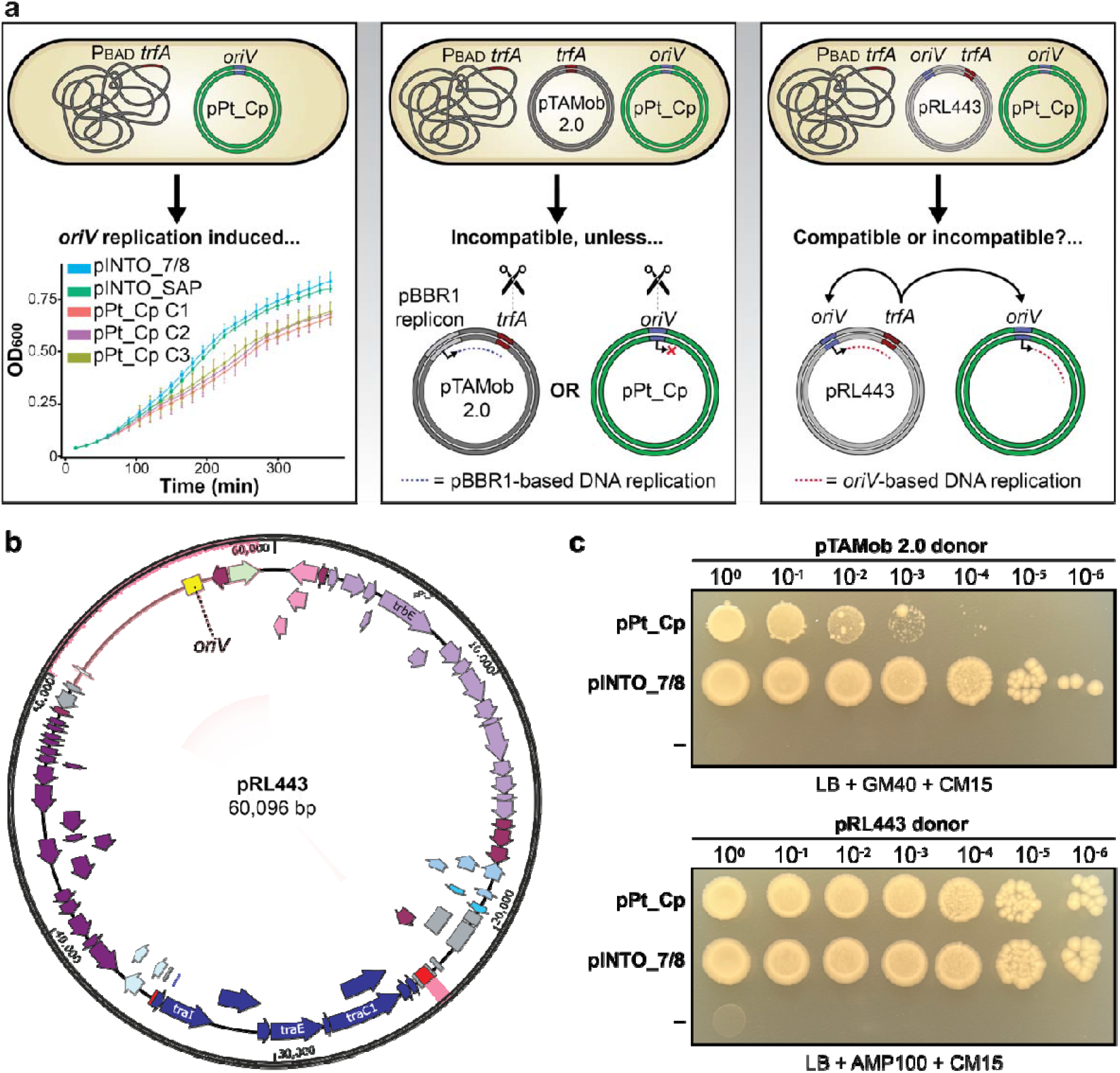
Conjugation using an alternative RK2 plasmid to further elucidate the cause of plasmid incompatibility. (**a**) <u>Panel 1</u>: past work by Walker et al. (2024) demonstrated that high-copy number *oriV* replication of pPt_Cp can be driven by inducing P_BAD_ *trfA* expression in *EPI300 E. coli*. This leads to a slight growth burden in the respective pPt_Cp clones (C1 to C3) when compared to EPI300 strains harbouring pCC1BAC-based cloning vectors (pINTO_7/8 and SAP). <u>Panel 2</u>: the plasmids pTAMob 2.0 and pPt_Cp cannot be co-maintained unless *trfA* is knocked-out or the *oriV* sequence is disrupted through indels. The conjugative plasmid pTAMob 2.0 has a pBBR1 replicon that drives its replication. <u>Panel 3</u>: to see if the pBBR1 replicon influences incompatibility, we sought to create a strain with pPt_Cp and pRL443, an RK2-based conjugative plasmid that replicates through the canonical RK2 *oriV/trfA* system. If pRL443 and pPt_Cp are compatible, this would suggest that the pBBR1 replicon is a driver of plasmid incompatibility. (**b**) A plasmid map of pRL443. The highlighted regions demonstrate portions of the plasmid that differ from pTAMob 2.0. This includes the RK2 replicon region (positioned at 50 to 59 kb) and the *aphA* kanamycin marker (positioned at 23 kb). (**c**) A spot plating assay where pTAMob 2.0 or pRL443 (i.e., donor strains) were conjugated into strains harbouring pPt_Cp or pINTO_7/8 (i.e., recipient strains). The negative control consists of the conjugative donor by itself. The antibiotic selection parameters are listed below the spot-plating images.

These observations led us to hypothesize that the pBBR1-based replicon in pTAMob 2.0 was initiating DNA replication at putative origin sites within the chloroplast genome (Fig. 5A, second panel), and that the resulting plasmid instability arose from the presence of multiple, competing replication origins dispersed throughout pPt_Cp. The presence of multiple replication origins on a single plasmid is known to disrupt replication control mechanisms, leading to reduced stability and inconsistent copy number. To test this, we performed conjugation using a strain harbouring pRL443 (Elhai et al., 1997), an RK2-based plasmid that employs the canonical *oriV*-TrfA replication system (Fig. 5A, third panel).

The conjugative plasmid pRL443 shares 87% sequence similarity to pTAMob 2.0 when globally aligned via EMBOSS Stretcher (Madeira et al., 2024), with there being two regions that could not be aligned (Fig. 5B). The first region spans ∼9.5 kb and includes the ampicillin and tetracycline resistance markers as well as the *oriV* replication origin, all of which are absent in pTAMob 2.0. The second region spans a mutated kanamycin resistance marker (*aphA*), which is disrupted by the URA3 yeast selection marker in pTAMob 2.0. Thus, the bulk of conjugative, partitioning, and regulatory genes are the same between both plasmids. When conjugated into the strains harbouring pPt_Cp or the cloning vector pINTO_7/8, pRL443 does not demonstrate any plasmid incompatibility, providing further evidence that the issue lies within the pBBR1 replicon in pTAMob 2.0.

The pBBR1 replicon was cloned in place of the RK2 *oriV* region during the construction of pTAMob to expand the host range of the conjugative plasmid (Strand et al., 2014). This replicon contains a putative origin of replication (*ori*) and replication initiator protein (RepB) that is thought to facilitate plasmid propagation in a range of gram-negative bacterial hosts (Kovach et al., 1995; Strand et al., 2014). Because pTAMob 2.0 only demonstrates incompatibility with pHALF_1/4, pTHIRDS_1/6, and pPt_Cp (Fig. 1B), we hypothesize that RepB recognizes and initiates replication at cryptic or putative origins present within the chloroplast genome.

### Analysis of the pBBR1-based replicon

The origin of replication (*ori*) for the pBBR1 replicon remains poorly characterized in the literature. To investigate its features, we examined a 2113 bp region of pTAMob 2.0 encompassing the annotated *repB* open reading frame. This region lies between the gentamicin resistance cassette and the yeast CEN/ARS/HIS sequences. Analysis with OriV-Finder (Li and Gao, 2025) confirmed the presence of the *repB* ORF and predicted an upstream 1005 bp region as the putative origin of replication, containing an AT-rich region, IHF binding motif, and three 18 bp iteron motifs (Supplemental Data File). In iteron-based plasmid replication systems, replication initiator proteins recognize and bind specific iteron sequences to trigger DNA replication. Accordingly, the putative pBBR1 iterons represent logical targets for probing potential origin-like regions within the chloroplast genome.

To characterize conserved features among the predicted iteron sequences, the three 18 bp motifs identified by OriV-Finder were aligned using WebLogo (Supplemental Fig. S4; Crooks et al., 2004). This analysis revealed a conserved core motif of NN<u>CGCAA</u>NNNN<u>T</u>N<u>ATTG</u>, highlighting sequence elements that may be important for RepB recognition and thus origin function. The consensus iteron sequence was then used to search the cloned *P. tricornutum* chloroplast genome using fuzznuc (Blankenberg et al., 2007; Rice et al., 2000), allowing up to one mismatch. This search identified seven dispersed matches across the 132.9 kb construct, none of which were localized to a specific region. The scattered distribution and lack of sites suggest that if iteron-like elements exist within the chloroplast genome, they are likely degenerate or functionally distinct from those associated with the pBBR1 replicon.

We also performed a local alignment of the 1005 bp putative pBBR1 *ori* to the cloned *P. tricornutum* chloroplast genome using EMBOSS Water (Rice et al., 2000), which identified a region of partial similarity spanning 1770 bp (37.5% identity, 52% gaps). The aligned region falls within the *rnl* (tRNA) locus of the inverted repeats. Interestingly, replication origins in the tobacco chloroplast genome have been mapped to the inverted repeats (Kunnimalaiyaan et al., 1997; K. Mühlbauer et al., 2002); however, replication can still proceed when these origins are inactivated, suggesting that multiple, possibly redundant, origins exist throughout the chloroplast genome (Scharff and Koop, 2007). Given the low overall similarity and extensive gaps, this alignment likely reflects coincidental sequence similarity rather than a putative pBBR1-like origin in the chloroplast genome.

## DISCUSSION

Conjugation-based methods offer a unique alternative for the delivery of partial or whole chloroplast genomes. Prior to our work, electroporation and chemical transformation methods had been used to clone chloroplast genomes in *E. coli* and yeast (Mordaka et al., 2025; O’Neill et al., 2012; Walker et al., 2024), and biolistic transformation had been used to deliver and integrate engineered chloroplast sequences into the endogenous chloroplast genome of *C. reinhardtii* (Mordaka et al., 2025; O’Neill et al., 2012). These methods all relied upon the *in vitro* isolation of DNA followed by shear physical forces to introduce chloroplast-derived constructs into the cell; comparatively, conjugation is an *in vivo* process that transfers mobilizable DNA between cells and is thus often more efficient for the movement of large (i.e., >50 kb) constructs. This work provides the first detailed exploration of bacterial conjugation as a strategy for transferring the chloroplast genome and identifies key factors governing compatibility between helper plasmids and chloroplast constructs.

We began our studies by investigating the source of incompatibility between pPt_Cp, the cloned *P. tricornutum* chloroplast genome, and pTAMob 2.0, a self-transmissible broad-host range conjugative plasmid derived from pTAMob (Strand et al., 2014). Disruption of the *oriV* sequence on pPt_Cp or knock-out of TrfA, the corresponding replication initiator protein present in pTAMob 2.0, restored plasmid compatibility. Interestingly, pPt_Cp can be replicated through the *oriV/trfA* system when maintained with pRL443, an alternative RK2-based plasmid, or when propagated by itself in the EPI300 *E. coli* strain, which contains an inducible genomic copy of *trfA*. Furthermore, when the chloroplast genome is cloned into smaller regions, plasmid incompatibility arises only in constructs containing half or more of the genome, suggesting that multiple disparate locations in the genome are interacting synergistically to drive plasmid incompatibility.

Taken together, our data suggests that the pBBR1 replicon present in pTAMob 2.0 is interacting with putative origins stippled throughout the chloroplast genome, and that competition for replication machinery between the TrfA/*oriV* origin and RepB/pBBR1-like origins is ultimately causing the emergence of plasmid instability. We sought to better characterize the pBBR1 origin using OriV-Finder (Li and Gao, 2025), which identified a 1005 bp sequence upstream of repB, the pBBR1 replication initiator protein. The putative iteron sequences and entire replication origin were subsequently analyzed to explore potential sequence features and homologies that may be present in the *P. tricornutum* chloroplast genome. Ultimately, no strong pBBR1-like iteron signatures or origins were identified within the chloroplast genome, suggesting that sequence homology alone is unlikely to account for the observed plasmid incompatibility.

Beyond plasmid compatibility, a method for direct conjugation to the chloroplast has yet to be developed. Conceptual approaches have been proposed in the literature, including the use of intracytoplasmic bacteria to directly conjugate to the chloroplast (Lim et al., 2008) and the addition of localization signals to conjugative proteins, like relaxase, to direct plasmid delivery to the chloroplast (Matsuoka and Maliga, 2021). An alternative would be to use the canonical conjugation method – that is, delivery to the nucleus – and then facilitate lateral plasmid transfer between the nucleus and chloroplast. Organelle-to-nucleus gene transfer (GT) has occurred extensively throughout evolutionary history, as evidenced by the thousands of nuclear-encoded genes that were once derived from the chloroplast and mitochondrial genomes (Kleine et al., 2009), but the opposite is not true. Nucleus-to-organelle GT appears to be exceedingly rare in nature, with there being a paucity of literature exploring this phenomenon in an experimental setting (Thorsness and Fox, 1990).

In conclusion, we have established an *E. coli* strain capable of efficiently mobilizing the *P. tricornutum* chloroplast genome. Our findings underscore the importance of carefully selecting and testing multiple helper plasmids when mobilizing large constructs like organelle genomes, as unanticipated incompatibilities can significantly hinder successful conjugation and thus downstream applications. This represents a promising alternative strategy for reintroducing cloned organelle genomes into algal cells, though further development is needed to make this approach practical. Overall, this study marks the first demonstration of conjugation as a means for mobilizing entire organelle genomes.

## METHODS

### Culturing conditions for E. coli, P. tricornutum, and S. cerevisiae

*E. coli* strain EPI300 (LGC Biosearch Technologies, Lucigen, catalog number: EC300110) was grown in lysogeny broth (LB, Miller formula) with or without 1.5% agar (w/v). Depending on the plasmid(s), LB was supplemented with 15 µg/ml chloramphenicol, 40 µg/ml gentamycin, 100 µg/ml ampicillin, and/or 100 µg/ml L-arabinose. *E. coli* strains were grown at 37ºC with liquid cultures being placed in a shaking incubator set to 225 rotations per minute (rpm).

*S. cerevisiae* strain VL6-48 (American Type Culture Collection [ATCC], catalog number: MYA-3666) was grown in 2X YPAD liquid media. Following yeast assembly, transformants were plated on complete minimal media lacking histidine (-HIS) or histidine and uracil (-HIS/URA) with 2% agar (w/v) and 1 M D-sorbitol. Yeast cultures were incubated at 30ºC with liquid cultures being placed in a shaking incubator set to 225 rotations per minute (rpm). Yeast YPAD and drop-out media were prepared as previously described (Walker et al., 2024).

### PCR amplification and preparation of fragments for assembly

Yeast assembly fragments were PCR amplified with GXL polymerase (Takara, catalog number: R050A) using the rapid protocol for amplicons larger than 5 kb or the slow protocol for all amplicons smaller than this (all primers are listed in Supplemental Table S3). Where applicable, optimized primers were generated using Primer3 (Untergasser et al., 2012). PCR-amplified fragments (50 to 100 µl) were treated with 10 units (0.5 µl) of *Dpn*I (New England Biolabs, catalog number: R0176) and incubated for 60 minutes at 37ºC followed by an enzyme de-activation step at 80ºC for 20 minutes. Enzyme-treated fragments for yeast assembly were directly used without further column purification.

### Multiplex PCR screening for pPt_Cp

Multiplex PCR screening was performed with SuperPlex premix (Takara, catalog number: 638543) and Primer3-optimized primer pairs (16 primers in total; Supplemental Table S3). We performed 10 µl MPX reactions using 1 µl of 10-times diluted DNA from the *E. coli* colonies being screened. For all screens, the positive control consisted of ∼10 ng of purified pPt_Cp plasmid DNA and the negative control consisted of ddH_2_O.

### DNA isolation from E. coli and S. cerevisiae

DNA were isolated from both species using modified alkaline lysis protocols as previously described (Walker et al., 2024). To isolate high-quality and intact pPt_Cp plasmid DNA for sequencing, we used the QIAGEN Large-Construct Kit (catalog number: 12462) as previously described (Walker et al., 2024). Plasmids with a pCC1BAC-based backbone (i.e., pQUART, pHALF, pTHIRDS, and pPt_Cp) were induced to high-copy number replication prior to DNA isolation. These cultures were first grown overnight in 5 ml LB supplemented with 15 µg/ml chloramphenicol. The following morning, 1.5 ml of saturated culture was inoculated into ∼23.5 ml of LB supplemented with 15 µg/ml chloramphenicol and 100 µg/ml L-arabinose. The induced cultures were grown for 5 to 7 hours before proceeding with DNA isolation. For pPt_Cp isolations, the volume of culture used for DNA isolation was scaled up to 500 ml.

### Yeast assembly of plasmids containing chloroplast DNA

Plasmids were assembled in *S. cerevisiae* using the same methods that were previously used to capture the cloned chloroplast genome (i.e., pPt_Cp; Karas et al., 2014; Walker et al., 2024). Quartered (pQUART), halved (pHALF), and three-quarter (pTHIRDS) genomes were assembled using the pINTO cloning vector. Overlapping chloroplast fragments were amplified using high-molecular weight *P. tricornutum* genomic DNA or the pre-cloned fragments (i.e., pCHAP plasmids) as a template for PCR. The pINTO backbone was amplified using 80 bp primers that would add 40 bp of homologous sequences to the respective chloroplast fragments at its termini (Supplemental Table S3). It was also split into two pieces that overlap in the yeast HIS3 marker to reduce the appearance of false positive transformants following assembly.

### Conjugation between bacterial strains

*E. coli* donor and recipient strains were inoculated from glycerol stocks into 5 ml of LB supplemented with the appropriate antibiotic. Cultures were grown overnight at 37ºC and 225 rpm for at least 8 hours. The following morning, the optical density (OD_600_) was measured for each culture (typically around an OD_600_ of 2-3), and then the cultures were diluted to an OD_600_ of 0.1 in 50 ml of fresh LB supplemented with the appropriate antibiotic. The cultures were grown until an OD_600_ of 1.0 was reached (∼3 hours); then, 45 ml of culture was transferred to a 50 ml conical tube and placed on ice. Cells were pelleted by centrifugation at 3000 x G for 10 minutes at 4ºC. After the first spin, the supernatant was decanted, and the cells were spun for an additional two minutes. Any remaining supernatant was removed using a P1000 pipette tip, and the pellet was resuspended with 300 µl of ice-cold 10% glycerol. The total resuspended cell volume was adjusted to 500 µl and then partitioned into 100 µl aliquots in 1.5 ml microtubes. These aliquots were placed in a -80ºC 95% ethanol bath to flash freeze the cells and then stored at -80ºC until use.

To perform conjugation, donor and recipient strains were removed from -80ºC storage and thawed on ice for ten minutes. For each conjugation, 100 µl of donor cells was combined with 100 µl of recipient cells and mixed by pipetting up-and-down five times. The entire mixture (200 µl) was then spread onto a room temperature 1.5% LB agar plate (w/v). Once dried (∼10 minutes), plates were transferred to a 37ºC incubator for 90 minutes, during which conjugation between the donor and recipient strains occurs.

After 90 minutes had passed, the plates were removed from the incubator and scraped using 2 ml of sddH_2_O. The scraped cells (1 to 1.5 ml) were transferred to a 1.5 ml microtube on ice, and then 200 µl of this volume was used to prepare serial dilutions of 10^0^ to 10^-6^ in a 96-well plate using sddH_2_O. For each dilution, 100 µl of the cells were spread with 100 µl of sddH_2_O onto 1.5% LB plates (w/v) supplemented with the appropriate antibiotics. For the spot plating assays, 5 µl of each dilution was plated and then dried for ∼20 minutes. The transconjugant plates were incubated at 37ºC for two days before being pictured.

### Electroporation to E. coli

Electroporation to *E. coli* was performed as previously described using homemade EPI300 electrocompetent cells (Walker et al., 2024). To recover strains harbouring only pPt_Cp or pTAMob 2.0 from the big CFUs, electroporated cells were spread across two 1.5% LB agar (w/v) plates – one that was supplemented with 40 µg/ml gentamycin (to select for pTAMob 2.0), and one with 15 µg/ml chloramphenicol (to select for pPt_Cp). Transformants appeared on the plates within 24 hours of incubation at 37ºC. To ensure that the transformants only contained one plasmid, single colonies were repatched onto two plates – one that contained the same antibiotic as they were originally plated on, and one that contained the alternative antibiotic. None of the transformants demonstrated double resistance for chloramphenicol and gentamycin, indicating that single plasmid uptake had occurred. Four colonies were pooled for every transformation and then used in the conjugation assays.

### Analysis of the pBBR1 replicon

A 2113 bp region of pTAMob 2.0 encompassing the annotated *repB* open reading frame was examined to identify replication origin features. The sequence was analyzed using OriV-Finder (Li and Gao, 2025) to predict replication-associated elements. OriV-Finder identified *repB* and an upstream 1005 bp region containing an AT-rich region, an integration host factor (IHF) binding site, and three putative 18 bp iteron sequences.

To examine conserved sequence features among these predicted iterons, the three 18 bp motifs were aligned using WebLogo v3 (Crooks et al., 2004), generating a consensus sequence of NNCGCAANNNNTNATTG. This consensus was used to search the cloned *Phaeodactylum tricornutum* chloroplast genome for related sequences using fuzznuc (Blankenberg et al., 2007; Rice et al., 2000) with the following parameters: pattern = NNCGCAANNNNTNATTG, mismatch = 1, and search of both DNA strands. This tool was accessed using the Galaxy bioinformatics platform (The Galaxy Community, 2024).

To evaluate broader sequence similarity, local pairwise alignment of the 1005 bp putative pBBR1 origin was performed against the cloned 132.9 kb *P. tricornutum* chloroplast genome sequence using EMBOSS Water (Rice et al., 2000). Alignments were generated using the EDNAFULL substitution matrix with a gap opening penalty of 10.0 and a gap extension penalty of 0.5.

## ACKNOWLEDGEMENTS

This work was funded by the Advanced Research + Invention Agency (ARIA) through project code PROP-PR01-P003, as well as the Natural Sciences and Engineering Research Council of Canada (NSERC) through project code RGPIN-2025-05428, both of which were awarded to B.J.K.

## AUTHOR CONTRIBUTIONS

**Conceptualization:** EJLW and BJK; **Data curation:** EJLW; **Formal analysis:** EJLW, BJK; **Funding acquisition:** BJK; **Investigation:** EJLW, TJ, AK; **Methodology:** EJLW, BJK; **Project administration:** BJK; **Resources:** BJK, **Supervision:** EJLW, BJK; **Validation:** EJLW, TJ, AK; **Visualization:** EJLW; **Writing:** EJLW; **Writing – review & editing:** EJLW, BJK.

