## Supplementary data file for "Resolving replication incompatibility between chloroplast and conjugative plasmids in *E. coli*"

**Predicted features from the pBBR1 replicon region (identified with OriV-Finder)**

CGTTGCTGCTCCATAACATCAAACATCGACCCACGGCGTAACGCGCTTGCTGCTTGGATGCCCGAGGCATAGACTGTACAAAAAAACAGTCATAACAAGCCATGAAAACCGCCACTGCGCCGTTACCACCGCTGCGTTCGGTCAAGGTTCTGGACCAGTTGCGTGAGCGCATACGCTACTTGCATTACAGTTTACGAACCGAACAGGCTTATGTCAACTGGGTTCGTGCCTTCATCCGTTTCCACGGTGTGCGTCCATGGGCAAATATTATACGCAAGGCGACAAGGTGCTGATGCCGCTGGCGATTCAGGTTCATCATGCCGTTTGTGATGGCTTCCATGTCGGCAGGAATTCGAATTCATACCCACCGGCTCCAACTGCGCGGCCTGCGGCCTTGCCCCATCAATTTTTTTAATTTTCTCTGGGGAAAAGCCTCCGGCCTGCGGCCTGCGCGCTTCGCTTGCCGGTTGGACACCAAGTGGAAGGCGGGTCAAGGCTCGCGCAGCGACCGCGCAGCGGCTTGGCCTTGACGCGCCTGGAACGACCCAAGCCTATGCGAGTGGGGGCAGTCGAAGGGCGAAGCCCGCCCGCCTGCCCCCCGAGCCTCACGGCGGCGAGTGCGGGGGTTCCAAGGGGGCAGCGCCACCTTGGGCAAGGCCGAAGGCCGCGCAGTCGATCAACAAGCCCCGGAGGGGCCACTTTTTGCCGGAGGGGGAGCCGCGCCGAAGGCGTGGGGGAACCCCGCAGGGGTGCCCT**TCTTTGGGCACCAAAGAACTAGATATAGGGCGAAATGCGAAAGACTTAAAAATCAACAACTTAAAAAAGGG**GG**GTACGCAACAGCTCATTG**CGGCACC**CCCCGCAATAGCTCATTG**CGTAGGTTAAAGAAAATCTGTAATTGACTGCCACTT**TTACGCAACGCATAATTG**TTGTCGCGCTGCCGAAAAGTTGCAGCTGATTGCGCATGGTGCCGCAACCGTGCGGCACCCCTACCGCATGGAGATAAGCATGGCCACGCAGTCCAGAGAAATCGGCATTCAAGCCAAGAACAAGCCCGGTCACTGGGTGCAAACGGAACGCAAAGCGCATGAGGCGTGGGCCGGGCTTATTGCGAGGAAACCCACGGCGGCAATGCTGCTGCATCACCTCGTGGCGCAGATGGGCCACCAGAACGCCGTGGTGGTCAGCCAGAAGACACTTTCCAAGCTCATCGGACGTTCTTTGCGGACGGTCCAATACGCAGTCAAGGACTTGGTGGCCGAGCGCTGGATCTCCGTCGTGAAGCTCAACGGCCCCGGCACCGTGTCGGCCTACGTGGTCAATGACCGCGTGGCGTGGGGCCAGCCCCGCGACCAGTTGCGCCTGTCGGTGTTCAGTGCCGCCGTGGTGGTTGATCACGACGACCAGGACGAATCGCTGTTGGGGCATGGCGACCTGCGCCGCATCCCGACCCTGTATCCGGGCGAGCAGCAACTACCGACCGGCCCCGGCGAGGAGCCGCCCAGCCAGCCCGGCATTCCGGGCATGGAACCAGACCTGCCAGCCTTGACCGAAACGGAGGAATGGGAACGGCGCGGGCAGCAGCGCCTGCCGATGCCCGATGAGCCGTGTTTTCTGGACGATGGCGAGCCGTTGGAGCCGCCGACACGGGTCACGCTGCCGCGCCGGTAG

**LEGEND:** putative type-I origin, AT-rich region, Integration Host Factor (IHF) binding motif, **iterons**, *repB* ORF
