## Supplementary tables and figures for "Resolving replication incompatibility between chloroplast and conjugative plasmids in *E. coli*"

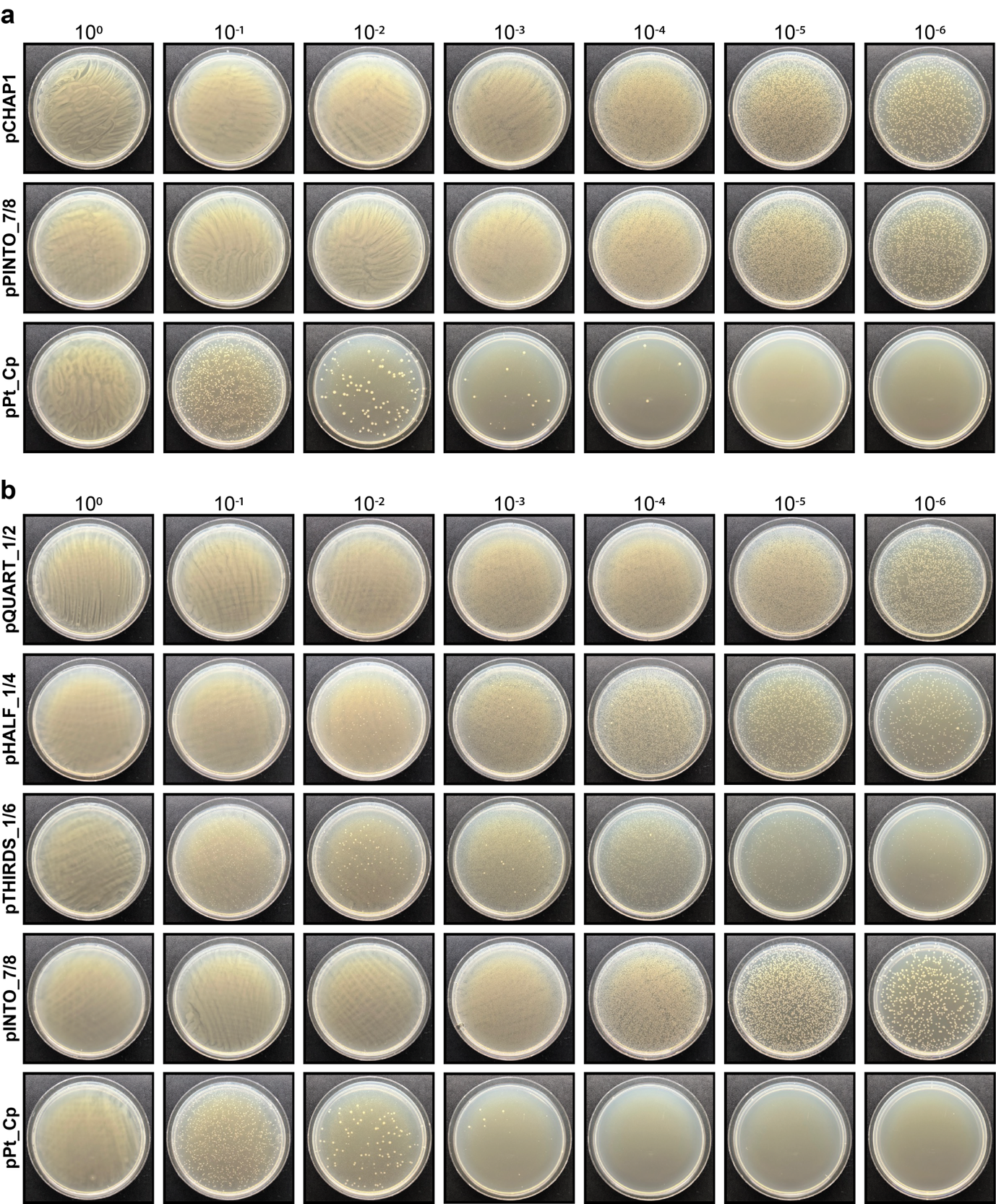

**Figure S1.** Plating following conjugation of pTAMob 2.0 into *E. coli* strains containing partial or complete plastid genomes. (**a**) Conjugation of pTAMob 2.0 into strains harbouring pCHAP1, pINTO_7/8, and pPt_Cp. (**b**) Conjugation of pTAMob 2.0 into strains harbouring pQUART_1/2, pHALF_1/4, pTHIRDS_1/6, pINTO_7/8, and pPt_Cp. Plating was performed on LB plates supplemented with chloramphenicol (15 µg/ml) and gentamycin (40 µg/ml).

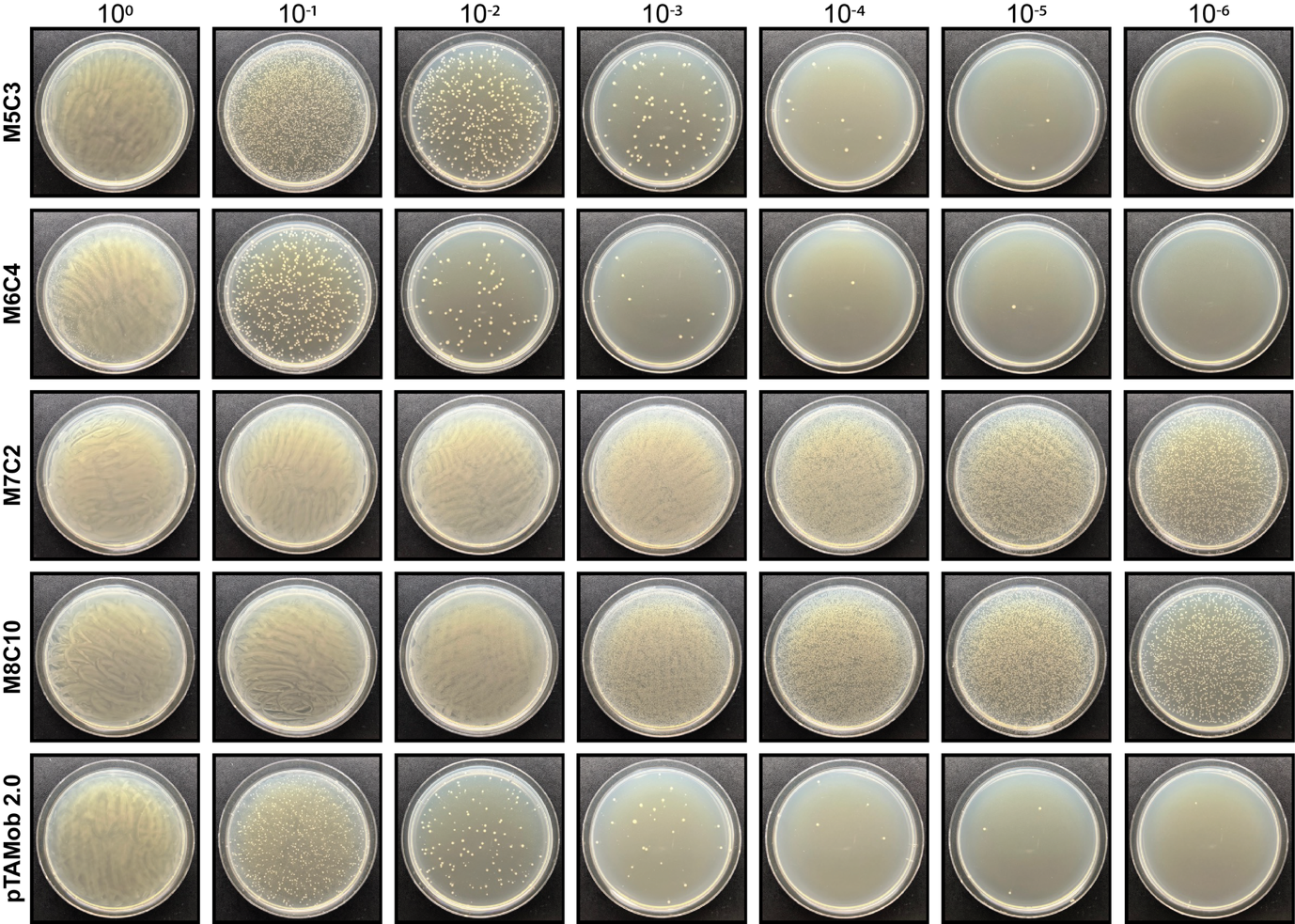

**Figure S2.** Plating following conjugation of minimized pTAMob 2.0 variants into an *E. coli* strain harbouring pPt_Cp*.* Plating was performed on LB plates supplemented with chloramphenicol (15 µg/ml) and gentamycin (40 µg/ml).

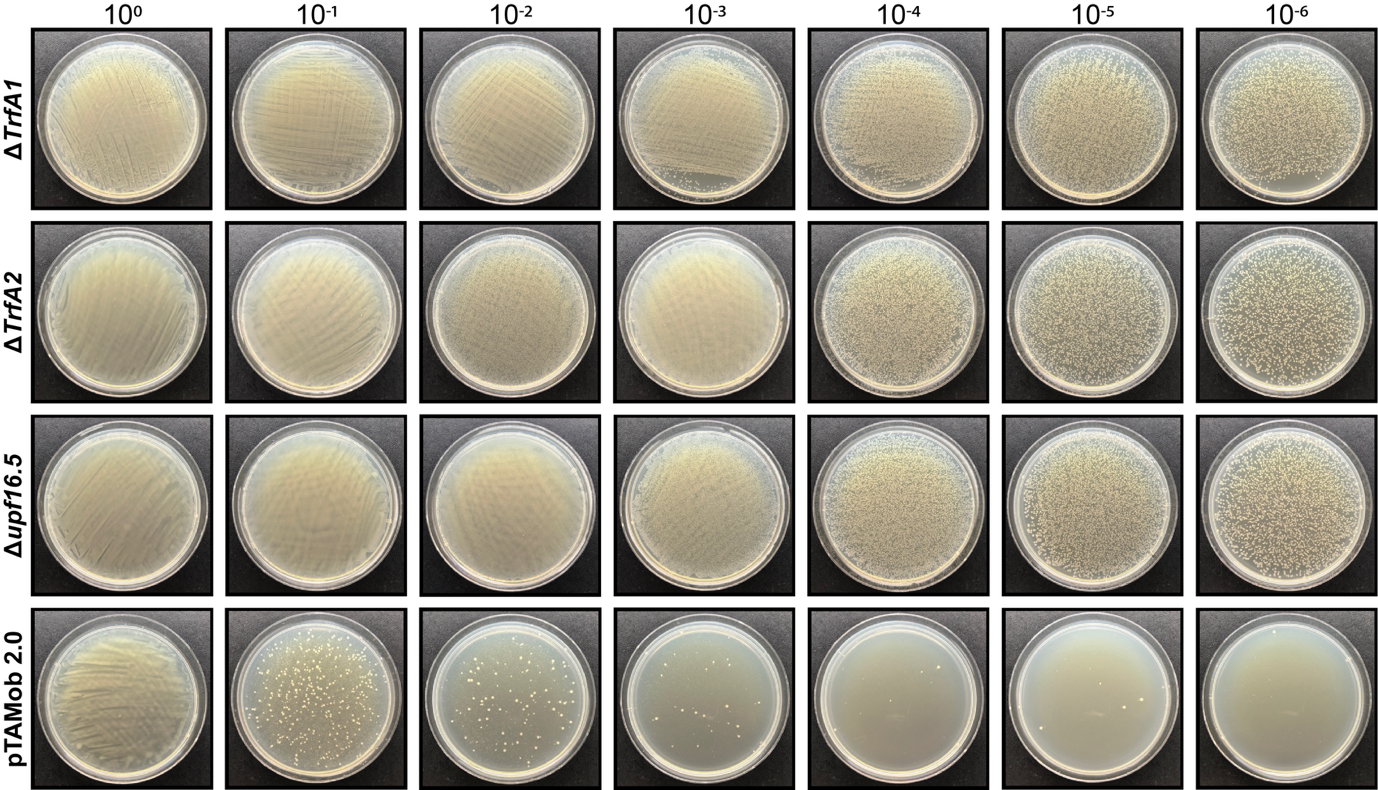

**Figure S3.** Plating following conjugation of deletion variants of pTAMob 2.0 into an *E. coli* strain harbouring pPt_Cp*.* Plating was performed on LB plates supplemented with chloramphenicol (15 µg/ml) and gentamycin (40 µg/ml).

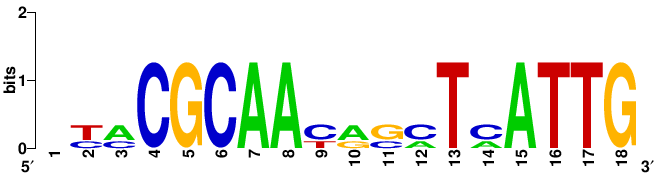

**Figure S4.** WebLogo visualization following multiple sequence alignment of three 18 bp putative iterons identified within the pBBR1 replicon using OriV-Finder. The consensus sequence NNCGCAANNNNTNATTG was derived from this alignment and represents conserved features potentially important for replication initiation.

**Table S1.** Addgene IDs for plasmids used in this study.

| **Plasmid** | **Addgene ID** |
| --- | --- |
| pCHAP1 | 206846 |
| pCHAP2 | 206847 |
| pCHAP3  a.k.a pCHAP3_SAP | 206849 |
| pCHAP4 | 206850 |
| pCHAP5 | 206851 |
| pCHAP6 | 206852 |
| pCHAP7 | 206853 |
| pCHAP8 | 206857 |
| pPt_Cp | 206855 |
| pSAP | 206429 |
| pINTO_7/8 | 206431 |
| pQUART_1/2 | 239653 |
| pQUART_3/4 | 239654 |
| pQUART_5/6 | 239655 |
| pQUART_7/8 | 239656 |
| pHALF_1/4 | 239657 |
| pHALF_5/8 | 239658 |
| pTHIRDS_1/6 | 239659 |
| pTAMob 2.0 | 149662 |
| M5C3 | 239660 |
| M6C4 | 239661 |
| M7C2 | 239662 |
| M8C10 | 239663 |
| pTAMob 2.0 Δ*trfA1* | Available upon request; not deposited to Addgene |
| pTAMob 2.0 Δ*trfA2* |  |
| pTAMob 2.0 Δupf16.5 |  |
| pRL443 | 70261 |

**Table S2**. Minimal variant (MV) regions identified in pTAMob 2.0. These regions were deleted in various combinations throughout minimal plasmids M5C3, M6C4, M7C2, and M8C10.

| **Region** | **Genes contained within region** |
| --- | --- |
| **MV6** | *trbN, trbO, trbP, upf31.7, fiwA, upf32.8, parA1, parA2, parB, parC, parD, parE, upf35.8, istA1, istA2, URA3, istB, upf35.8, aphA, traA, rtaB, traC1, traC2, traD, traE* |
| **MVA** | *klaC, klaB, klaA, kleF1, kleF2, kleE, kleD, kleC, kleB, kleA* |
| **MVC** | *traJ*, traX*, traI*, upf16.5, trfA1, trfA2* |

*Present in more than one copy in pTAMob 2.0

**Table S3.** Oligonucleotides used in this study.

| **Amplicon** | **Primers (5′ to 3′)** | **Template** | **Size (bp)** |
| --- | --- | --- | --- |
| **Assembly of pQUART_1/2 – selection on -HIS media** | | | |
| Chloroplast fragment 1 | **F**: tggtgaaaacgttgaaatcg  **R**: gtccaggtcgtggtggtact | Genomic *P. tricornutum* DNA | 14,543 |
| Chloroplast fragment 2 | **F**: acatccaagcccagactgat  **R**: ttgttttcgttggttggtca |  | 15,249 |
| Backbone fragment 1 | **F**:taataataaacctgaccaaccaacgaaaacaaTGAAGAGCATCTTCCGCTGCATAACCCTGCTTCGGGGTCATTATAGCG  **R**: TTCAGTGGTGTGATGGTCGT | pINTO_7/8 | 6503 |
| Backbone fragment 2 | **F**: CAGTAGCAGAACAGGCCACA  **R**:cggtaacacccacgatttcaacgttttcaccatGAAGAGCAAACCAAAGCGGAGTGACTGCAACTAATGAAAATTAAATT |  | 7994 |
| **Assembly of pQUART_3/4 – selection on -HIS media** | | | |
| Chloroplast fragment 3 | **F:** aagaccaccgatttgacacc  **R:**tgcacctgcgcgacatgaataaggtaattcaataccttgctcCtcagcagcatctaaaac | pCHAP3_SAP | 17,278 bp |
| Chloroplast fragment 4 | **F:**tgattgtaatgatgatgtatttgttttagatgctgctgaGgagcaaggtattgaattacc  **R:** CGCCATGTTCAGCAAGTAAA | Genomic *P. tricornutum* DNA |  |
| Backbone fragment 1 | **F**:TGAATTAAATATTTTACTTGCTGAACATGGCGTGAAGAGCATCTTCCGCTGCATAACCCTGCTTCGGGGTCATTATAGCG  **R**: TTCAGTGGTGTGATGGTCGT | pINTO_7/8 | 6503 |
| Backbone fragment 2 | **F:** CAGTAGCAGAACAGGCCACA  **R:**atttcacacgttggtgtcaaatcggtggtctttGAAGAGCAAACCAAAGCGGAGTGACTGCAACTAATGAAAATTAAATT |  | 7994 |
| **Assembly of pQUART_5/6 – selection on -HIS media** | | | |
| Chloroplast fragment 5 | **F**: ggtttaaaacattagtgggtgga  **R**: ccatgtgttgttccaacgag | Genomic *P. tricornutum* DNA | 17,639 |
| Chloroplast fragment 6 | **F**: ttgctcgggtcttagctgat  **R**: actggccggtttcattgtag |  | 15,671 |
| Backbone fragment 1 | **F**:ATATTGTTCAATCTACAATGAAACCGGCCAGTTGAAGAGCATCTTCCGCTGCATAACCCTGCTTCGGGGTCATTATAGCG  **R**: TTCAGTGGTGTGATGGTCGT | pINTO_7/8 | 6503 |
| Backbone fragment 2 | **F**: CAGTAGCAGAACAGGCCACA  **R**:GTAAACTGTTCCACCCACTAATGTTTTAAACCtGAAGAGCAAACCAAAGCGGAGTGACTGCAACTAATGAAAATTAAATT |  | 7994 |
| **Assembly of pQUART_7/8 – selection on -HIS media** | | | |
| Chloroplast fragment 7 | **F**: tggaatttagttgggttacgc  **R**: tcgtcggcaaaaaccttatc | Genomic *P. tricornutum* DNA | 14,092 |
| Chloroplast fragment 8 | **F**: acattcgcatgtcacctcat  **R**: ttatcaccggcaaaaccttc |  | 11,780 |
| Backbone fragment 1 | **F**:aaaactttagaggaaggttttgccggtgataaTGAAGAGCATCTTCCGCTGCATAACCCTGCTTCGGGGTCATTATAGCG  **R**: TTCAGTGGTGTGATGGTCGT | pINTO_7/8 | 6503 |
| Backbone fragment 2 | **F**: CAGTAGCAGAACAGGCCACA  **R**:attgttttaaagcgtaacccaactaaattccatGAAGAGCAAACCAAAGCGGAGTGACTGCAACTAATGAAAATTAAATT |  | 7994 |
| **Assembly of pHALF_1/4 – selection on -HIS/URA media** | | | |
| Chloroplast fragment 1 | **Same specifications as listed for pQUART_1/2 and 3/4** | | |
| Chloroplast fragment 2 |  |  |  |
| Chloroplast fragment 3 |  |  |  |
| Chloroplast fragment 4 |  |  |  |
| Backbone fragment 1 | **F:**TGAATTAAATATTTTACTTGCTGAACATGGCGTGAAGAGCATCTTCCGCTGCATAACCCTGCTTCGGGGTCATTATAGCG  **R:** TTCAGTGGTGTGATGGTCGT | pINTO_7/8 | 6503 |
| Backbone fragment 2 | **F:** CAGTAGCAGAACAGGCCACA  **R:**cggtaacacccacgatttcaacgttttcaccatGAAGAGCAAACCAAAGCGGAGTGACTGCAACTAATGAAAATTAAATT |  | 7994 |
| **Assembly of pHALF_5/8 – selection on -HIS media** | | | |
| Chloroplast fragment 5 | **Same specifications as listed for pQUART_5/6 and 7/8** | | |
| Chloroplast fragment 6 |  |  |  |
| Chloroplast fragment 7 |  |  |  |
| Chloroplast fragment 8 |  |  |  |
| Backbone fragment 1 | **F:**aaaactttagaggaaggttttgccggtgataaTGAAGAGCATCTTCCGCTGCATAACCCTGCTTCGGGGTCATTATAGCG  **R:** TTCAGTGGTGTGATGGTCGT | pINTO_7/8 | 6503 |
| Backbone fragment 2 | **F:** CAGTAGCAGAACAGGCCACA  **R:**GTAAACTGTTCCACCCACTAATGTTTTAAACCtGAAGAGCAAACCAAAGCGGAGTGACTGCAACTAATGAAAATTAAATT |  | 7994 |
| **Assembly of pTHIRDS_1/6 – selection on -HIS/URA media** | | | |
| Chloroplast fragment 1 | **Same specifications as listed for pQUART_1/2, 3/4, and 5/6** | | |
| Chloroplast fragment 2 |  |  |  |
| Chloroplast fragment 3 |  |  |  |
| Chloroplast fragment 4 |  |  |  |
| Chloroplast fragment 5 |  |  |  |
| Chloroplast fragment 6 |  |  |  |
| Backbone fragment 1 | **F**:ATATTGTTCAATCTACAATGAAACCGGCCAGTTGAAGAGCATCTTCCGCTGCATAACCCTGCTTCGGGGTCATTATAGCG  **R**: TTCAGTGGTGTGATGGTCGT | pINTO_7/8 | 6503 |
| Backbone fragment 2 | **F**: CAGTAGCAGAACAGGCCACA  **R**:cggtaacacccacgatttcaacgttttcaccatGAAGAGCAAACCAAAGCGGAGTGACTGCAACTAATGAAAATTAAATT |  | 7994 |
| **Multiplex screening primers for pPt_Cp** | | | |
| Amplicon in fragment 1 | **F**: agcaattcgacgtttacccg  **R**: ctaatggtgagcttgcgtcg | DNA isolated from pPt_Cp *E. coli* transconjugants or transformants | 650 |
| Amplicon in fragment 2 | **F**: cgctttatattggccgcctt  **R**: aaaacgcaaagagaagccgt |  | 504 |
| Amplicon in fragment 3 | **F**: ggggatcgaggaacaggaaa  **R**: ccttcgcgttcaactgtgtt |  | 404 |
| Amplicon in fragment 4 | **F**: aaatcatttccttcgcgcgt  **R**: ggtactccaacgttaggtcg |  | 329 |
| Amplicon in fragment 5 | **F**: gcgaaggaattggaagcgaa  **R**: gaccatgtactagaaatcctgca |  | 269 |
| Amplicon in fragment 6 | **F**: tgtctacaaaatgcgtccaaaga  **R**: aacctgttcaccaccaaagc |  | 213 |
| Amplicon in fragment 7 | **F**: aagaaggtatcgtgccaggt  **R**: accagcattttccgcaatcc |  | 161 |
| Amplicon in fragment 8 | **F**: ggtgatttagagggtagacggt  **R**: tcaattgctaggctatagttcca |  | 137 |
